## Supplementary Figures for "Identification of protein-protected mRNA fragments and structured excised intron RNAs in human plasma by TGIRT-seq peak calling"

**Fig. S1**

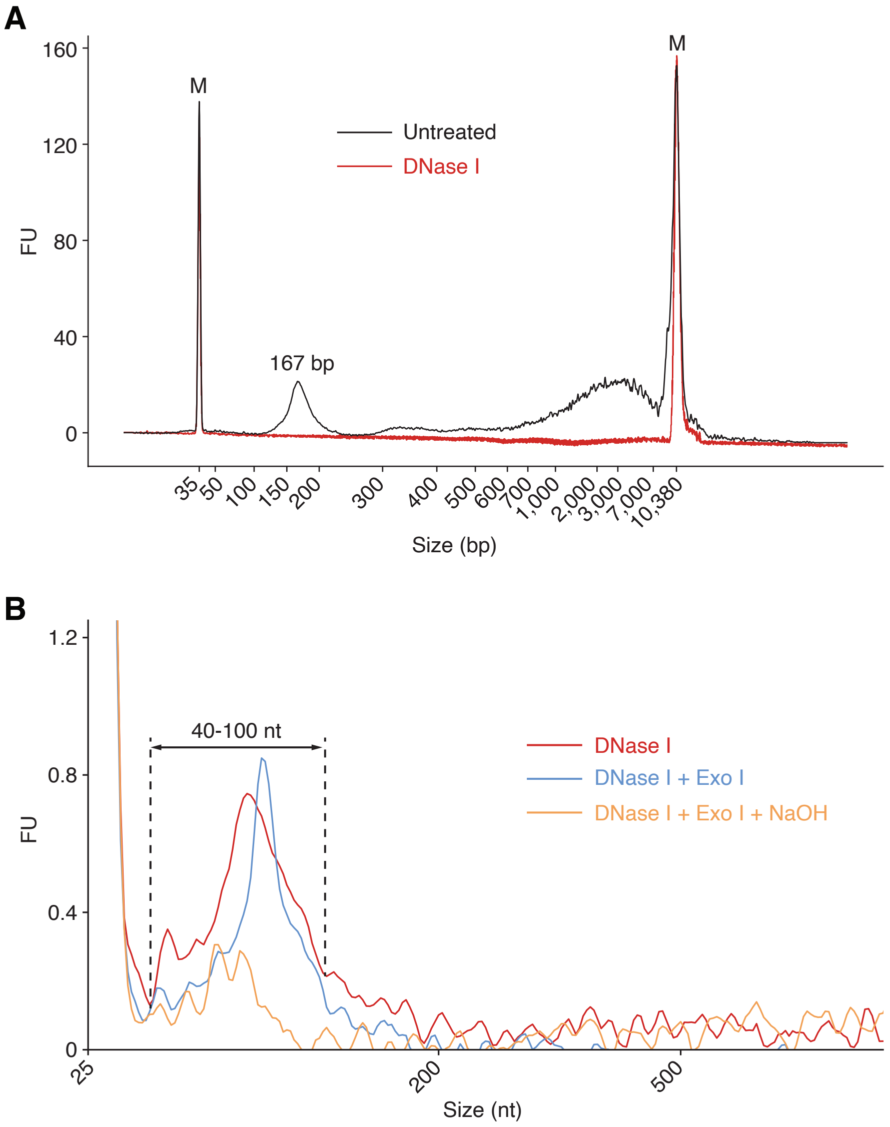
A
M
M
Untreated
DNase I
167 bp
FU
160
120
80
40
0
35
50
100
150
200
300
400
500
600
700
1,000
2,000
3,000
7,000
10,380
Size (bp)
B
40-100 nt
DNase I
DNase I + Exo I
DNase I + Exo I + NaOH
FU
1.2
0.8
0.4
0
25
200
500
Size (nt)

#### Fig. S2

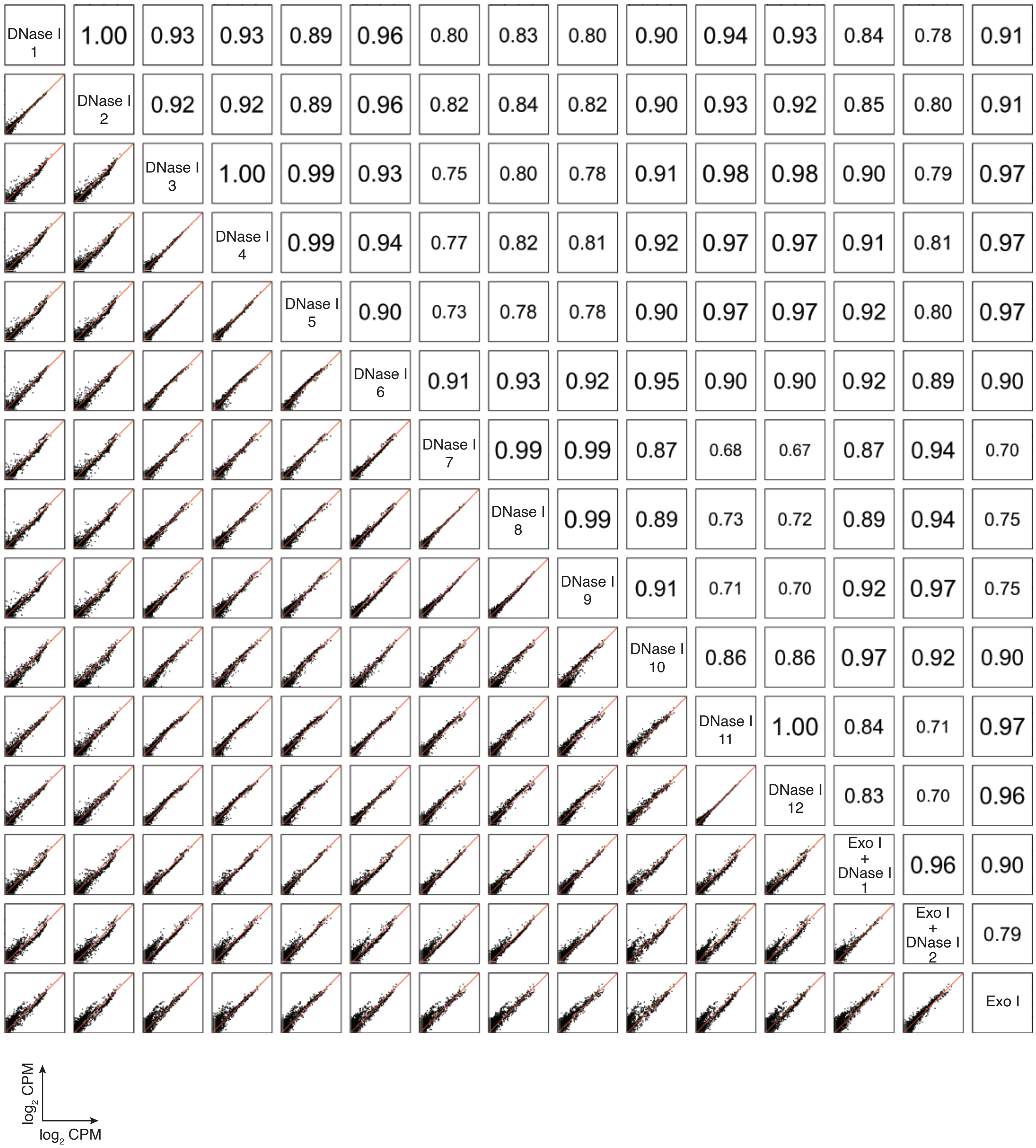
DNase I 1
1.00
0.93
0.93
0.89
0.96
0.80
0.83
0.80
0.90
0.94
0.93
0.84
0.78
0.91
DNase I 2
0.92
0.92
0.89
0.96
0.82
0.84
0.82
0.90
0.93
0.92
0.85
0.80
0.91
DNase I 3
1.00
0.99
0.93
0.75
0.80
0.78
0.91
0.98
0.98
0.90
0.79
0.97
DNase I 4
0.99
0.94
0.77
0.82
0.81
0.92
0.97
0.97
0.91
0.81
0.97
DNase I 5
0.90
0.73
0.78
0.78
0.90
0.97
0.97
0.92
0.80
0.97
DNase I 6
0.91
0.93
0.92
0.95
0.90
0.90
0.92
0.89
0.90
DNase I 7
0.99
0.99
0.87
0.68
0.67
0.87
0.94
0.70
DNase I 8
0.99
0.89
0.73
0.72
0.89
0.94
0.75
DNase I 9
0.91
0.71
0.70
0.92
0.97
0.75
DNase I 10
0.86
0.86
0.97
0.92
0.90
DNase I 11
1.00
0.84
0.71
0.97
DNase I 12
0.83
0.70
0.96
Exo I + DNase I 1
0.96
0.90
Exo I + DNase I 2
0.79
Exo I
log2 CPM
log2 CPM

#### Fig. S3

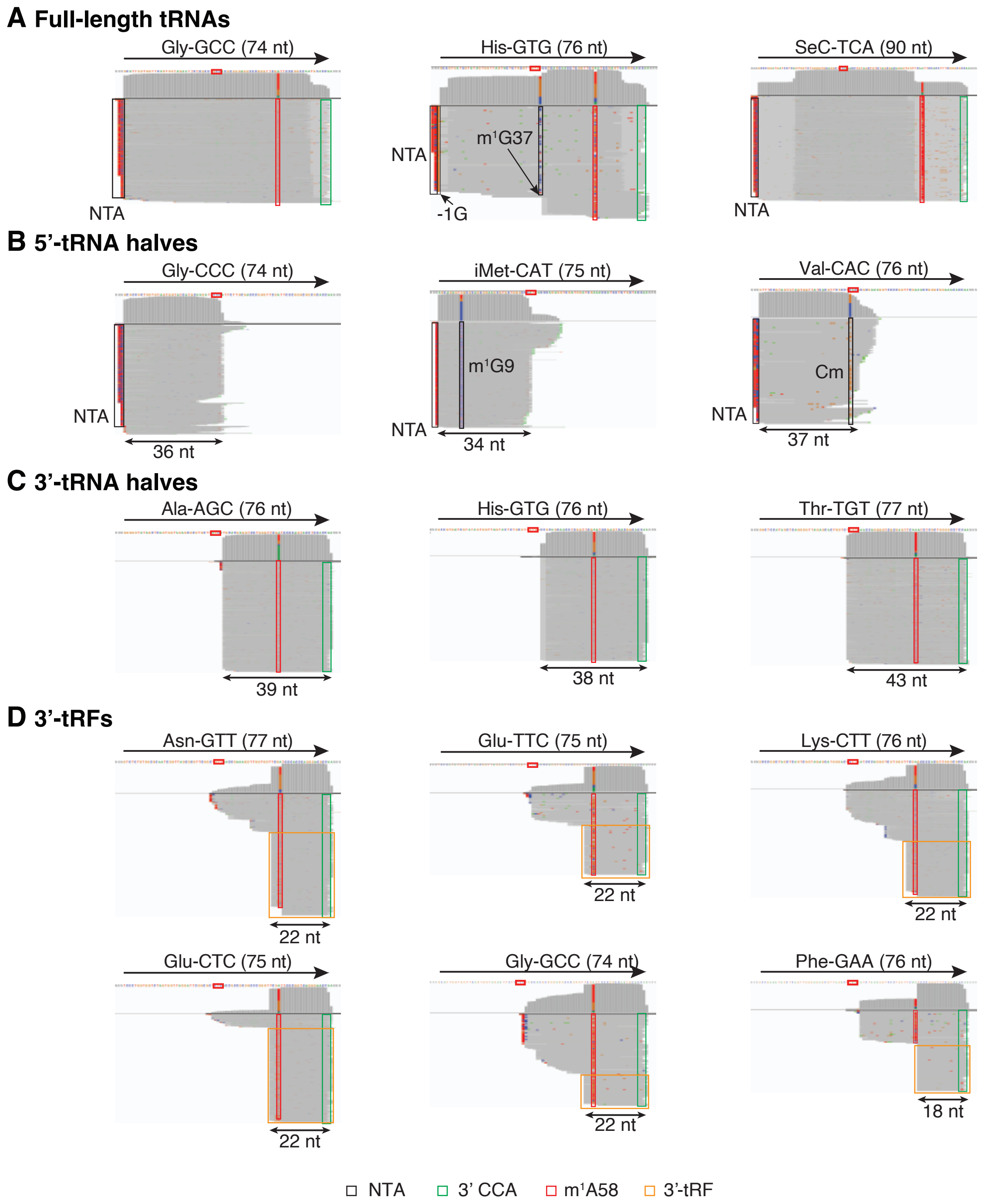

A Full-length tRNAs
Gly-GCC (74 nt)
NTA
His-GTG (76 nt)
NTA
m1G37
-1G
SeC-TCA (90 nt)
NTA
B 5'-tRNA halves
Gly-CCC (74 nt)
NTA
36 nt
iMet-CAT (75 nt)
m1G9
NTA
34 nt
Val-CAC (76 nt)
Cm
NTA
37 nt
C 3'-tRNA halves
Ala-AGC (76 nt)
39 nt
His-GTG (76 nt)
38 nt
Thr-TGT (77 nt)
43 nt
D 3'-tRFs
Asn-GTT (77 nt)
22 nt
Glu-TTC (75 nt)
22 nt
Lys-CTT (76 nt)
22 nt
Glu-CTC (75 nt)
22 nt
Gly-GCC (74 nt)
22 nt
Phe-GAA (76 nt)
18 nt
NTA
3' CCA
m1A58
3'-tRF

### Fig. S4

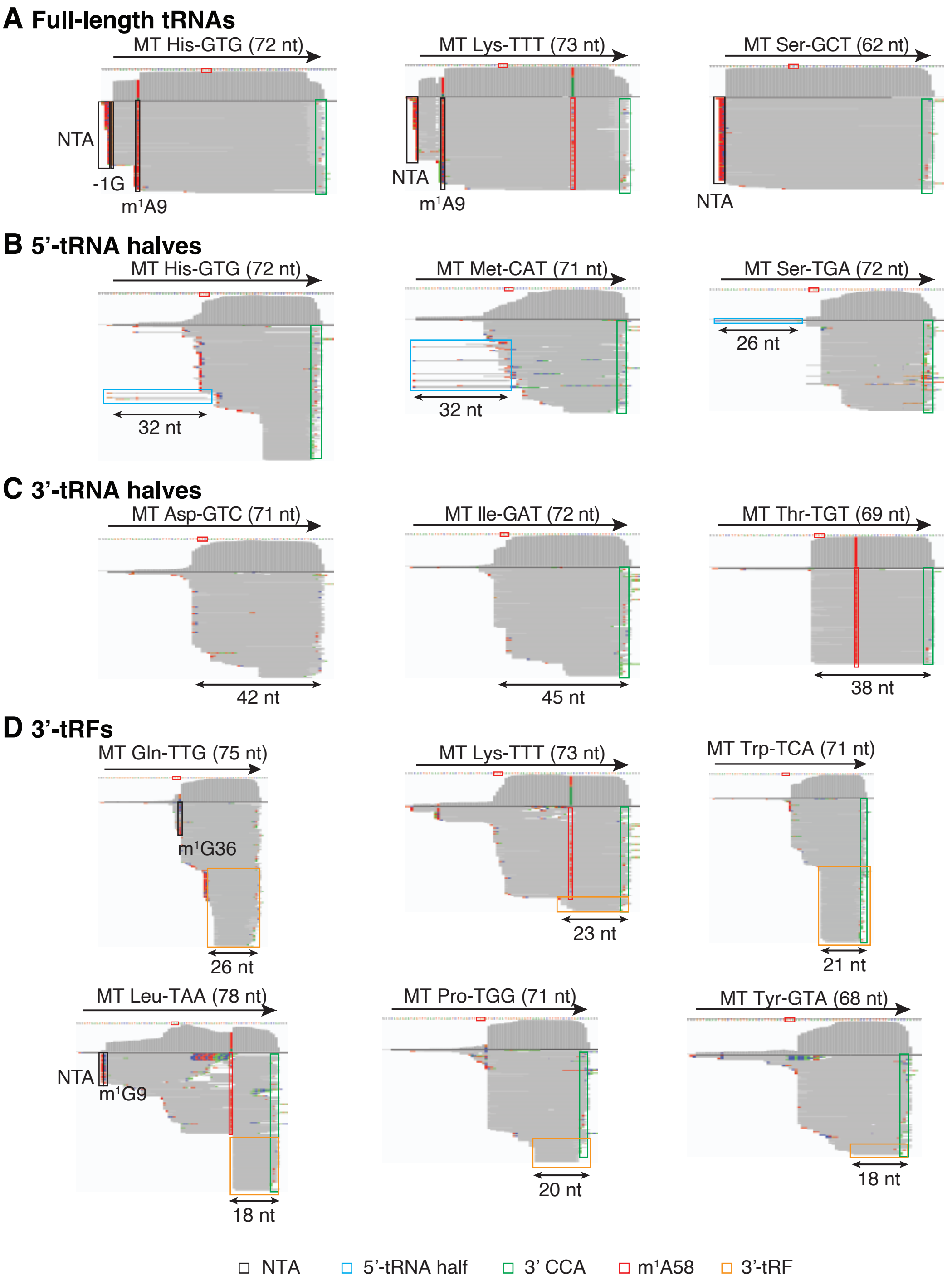

A Full-length tRNAs
MT His-GTG (72 nt)
MT Lys-TTT (73 nt)
MT Ser-GCT (62 nt)
NTA
-1G
m1A9
NTA
m1A9
NTA
B 5'-tRNA halves
MT His-GTG (72 nt)
MT Met-CAT (71 nt)
MT Ser-TGA (72 nt)
32 nt
32 nt
26 nt
C 3'-tRNA halves
MT Asp-GTC (71 nt)
MT Ile-GAT (72 nt)
MT Thr-TGT (69 nt)
42 nt
45 nt
38 nt
D 3'-tRFs
MT Gln-TTG (75 nt)
MT Lys-TTT (73 nt)
MT Trp-TCA (71 nt)
m1G36
26 nt
23 nt
21 nt
MT Leu-TAA (78 nt)
MT Pro-TGG (71 nt)
MT Tyr-GTA (68 nt)
NTA
m1G9
18 nt
20 nt
18 nt
NTA
5'-tRNA half
3' CCA
m1A58
3'-tRF

#### Fig. S5

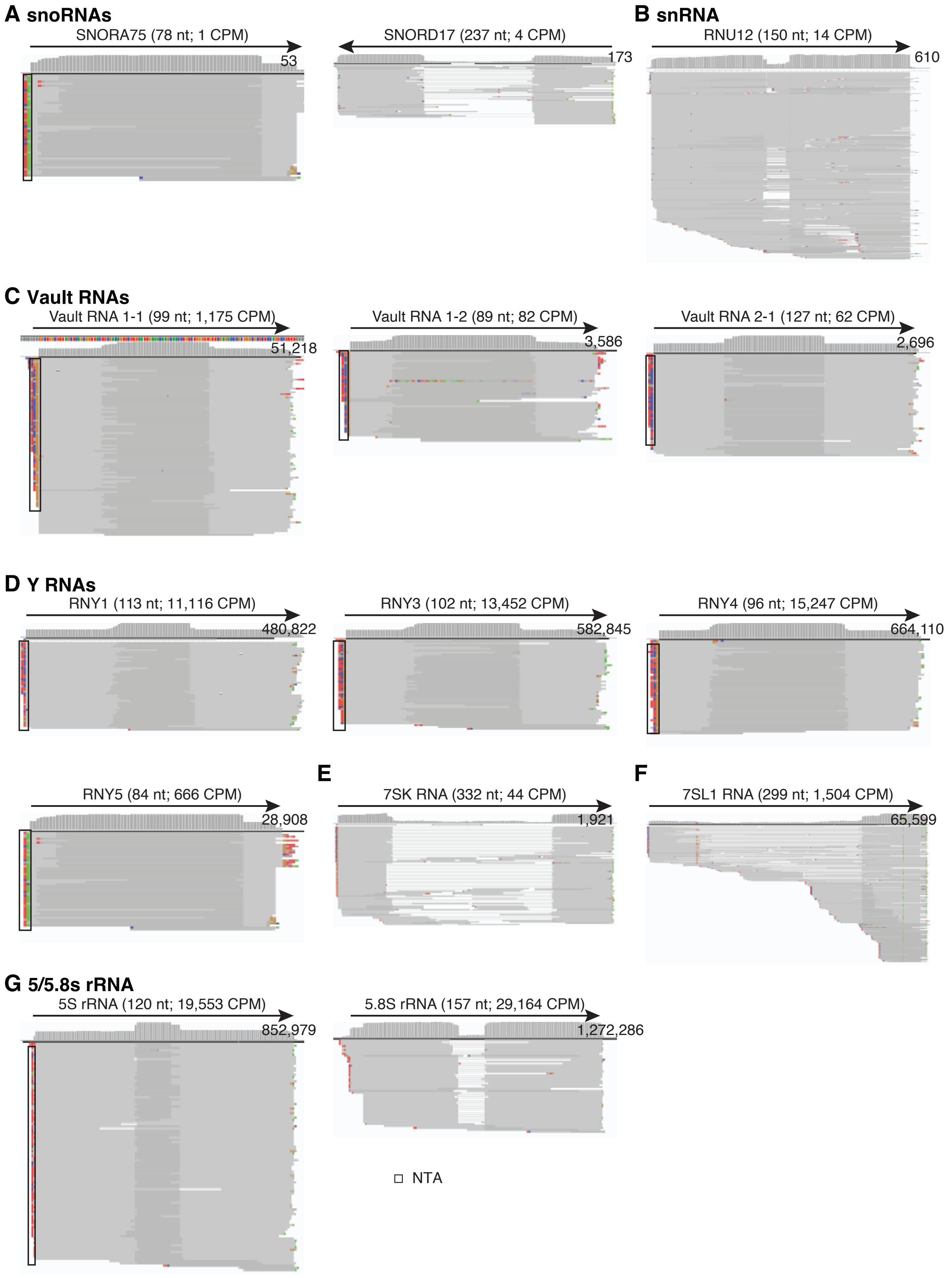
A snoRNAs
SNORA75 (78 nt; 1 CPM)
53
SNORD17 (237 nt; 4 CPM)
173
B snRNA
RNU12 (150 nt; 14 CPM)
610
C Vault RNAs
Vault RNA 1-1 (99 nt; 1,175 CPM)
51,218
Vault RNA 1-2 (89 nt; 82 CPM)
3,586
Vault RNA 2-1 (127 nt; 62 CPM)
2,696
D Y RNAs
RNY1 (113 nt; 11,116 CPM)
480,822
RNY3 (102 nt; 13,452 CPM)
582,845
RNY4 (96 nt; 15,247 CPM)
664,110
RNY5 (84 nt; 666 CPM)
28,908
E
7SK RNA (332 nt; 44 CPM)
1,921
F
7SL1 RNA (299 nt; 1,504 CPM)
65,599
G 5/5.8s rRNA
5S rRNA (120 nt; 19,553 CPM)
852,979
5.8S rRNA (157 nt; 29,164 CPM)
1,272,286
NTA

#### Fig. S6

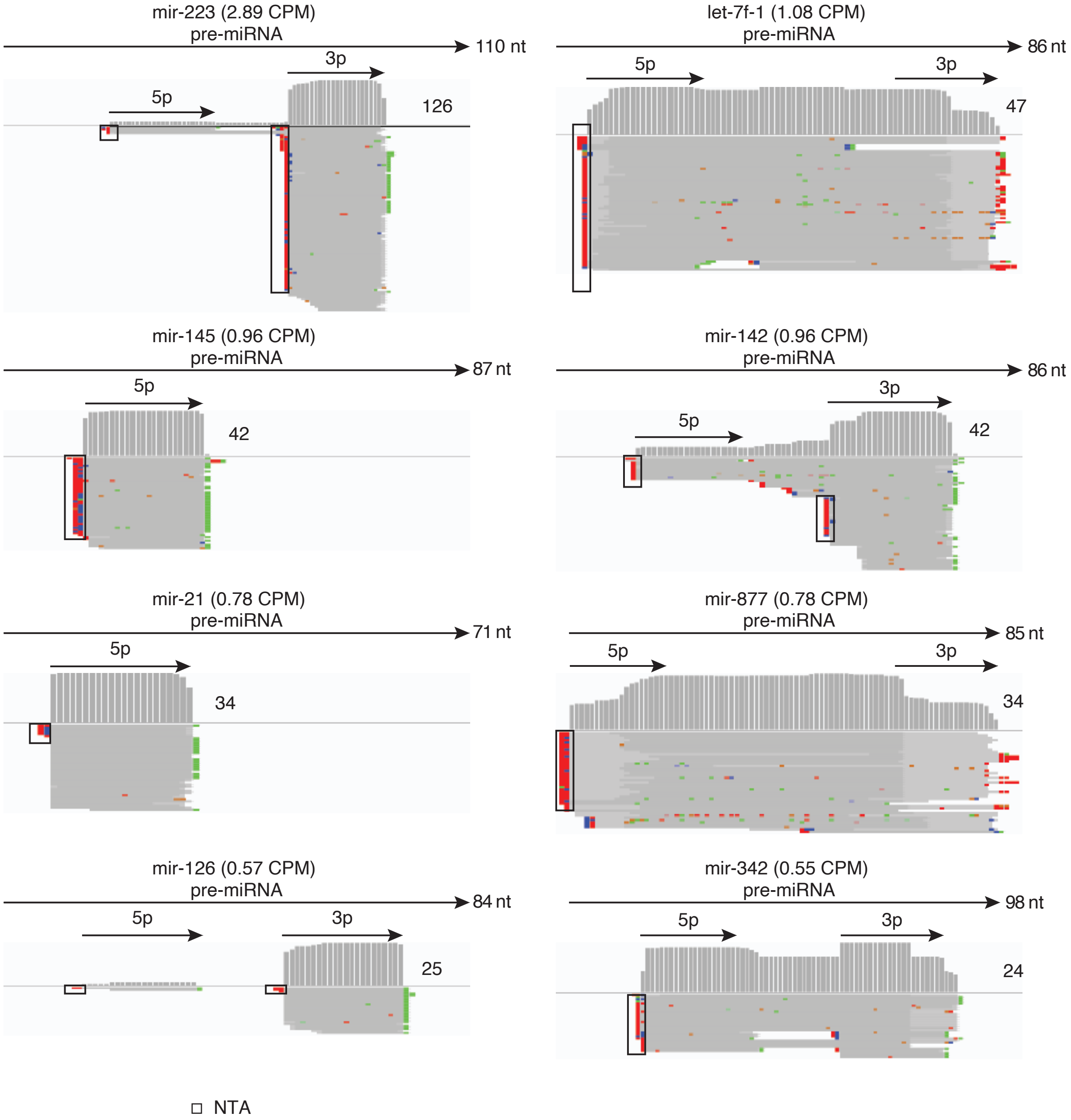

mir-223 (2.89 CPM)
pre-miRNA
110 nt
5p
3p
126
let-7f-1 (1.08 CPM)
pre-miRNA
86 nt
5p
3p
47
mir-145 (0.96 CPM)
pre-miRNA
87 nt
5p
42
mir-142 (0.96 CPM)
pre-miRNA
86 nt
5p
3p
42
mir-21 (0.78 CPM)
pre-miRNA
71 nt
5p
34
mir-877 (0.78 CPM)
pre-miRNA
85 nt
5p
3p
34
mir-126 (0.57 CPM)
pre-miRNA
84 nt
5p
3p
25
mir-342 (0.55 CPM)
pre-miRNA
98 nt
5p
3p
24
NTA

**Fig. S7**

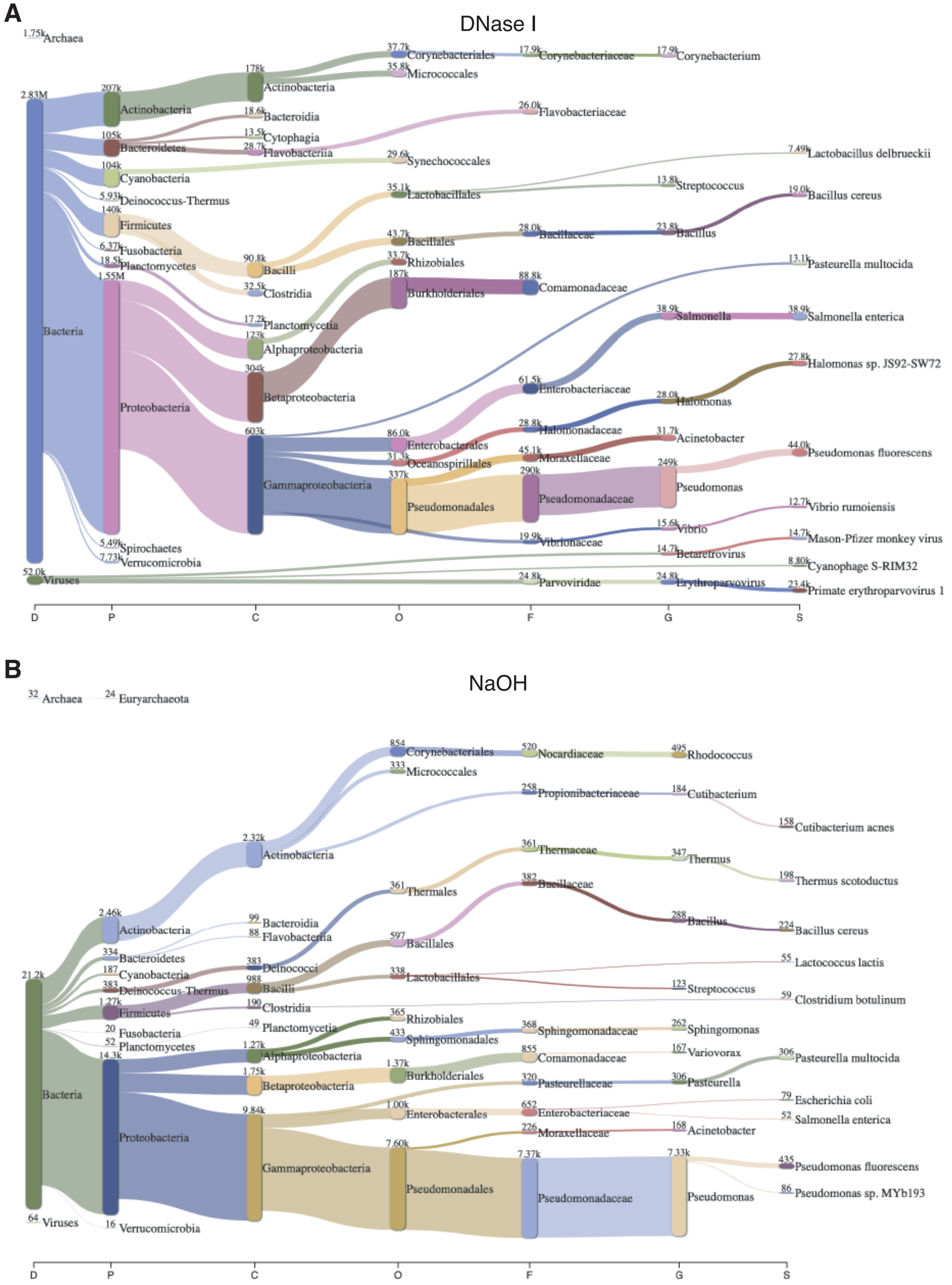

A
DNase I
1.75k Archaea
2.83M
Bacteria
207k Actinobacteria
178k Actinobacteria
37.7k Corynebacteriales
17.9k Corynebacteriaceae
17.9k Corynebacterium
35.8k Micrococcales
105k Bacteroidetes
18.6k Bacteroidia
13.5k Cytophagia
28.7k Flavobacteriia
26.0k Flavobacteriaceae
104k Cyanobacteria
29.6k Synechococcales
5.93k Deinococcus-Thermus
140k Firmicutes
35.1k Lactobacillales
7.49k Lactobacillus delbrueckii
13.8k Streptococcus
19.0k Bacillus cereus
43.7k Bacillales
28.0k Bacillaceae
23.8k Bacillus
6.37k Fusobacteria
18.5k Planctomycetes
90.8k Bacilli
32.5k Clostridia
33.7k Rhizobiales
13.1k Pasteurella multocida
1.55M Proteobacteria
187k Burkholderiales
88.8k Comamonadaceae
17.2k Planctomycetia
123k Alphaproteobacteria
38.9k Salmonella
38.9k Salmonella enterica
304k Betaproteobacteria
27.8k Halomonas sp. JS92-SW72
61.5k Enterobacteriaceae
28.0k Halomonas
603k Gammaproteobacteria
86.0k Enterobacterales
28.8k Halomonadaceae
31.7k Acinetobacter
31.3k Oceanospirillales
45.1k Moraxellaceae
44.0k Pseudomonas fluorescens
337k Pseudomonadales
290k Pseudomonadaceae
249k Pseudomonas
12.7k Vibrio rumoiensis
15.6k Vibrio
19.9k Vibrionaceae
14.7k Mason-Pfizer monkey virus
5.49k Spirochaetes
7.73k Verrucomicrobia
14.7k Betaretrovirus
8.80k Cyanophage S-RIM32
52.0k Viruses
24.8k Parvoviridae
24.8k Erythroparvovirus
23.4k Primate erythroparvovirus 1
D
P
C
O
F
G
S
B
NaOH
32 Archaea
24 Euryarchaeota
854 Corynebacteriales
520 Nocardiaceae
495 Rhodococcus
333 Micrococcales
258 Propionibacteriaceae
184 Cutibacterium
158 Cutibacterium acnes
2.32k Actinobacteria
361 Thermaceae
347 Thermus
198 Thermus scotoductus
361 Thermales
382 Bacillaceae
288 Bacillus
224 Bacillus cereus
2.46k Actinobacteria
99 Bacteroidia
88 Flavobacteriia
597 Bacillales
334 Bacteroidetes
383 Deinococci
55 Lactococcus lactis
187 Cyanobacteria
988 Bacilli
338 Lactobacillales
123 Streptococcus
21.2k
383 Deinococcus-Thermus
59 Clostridium botulinum
1.27k Firmicutes
190 Clostridia
365 Rhizobiales
20 Fusobacteria
49 Planctomycetia
433 Sphingomonadales
368 Sphingomonadaceae
262 Sphingomonas
52 Planctomycetes
855 Comamonadaceae
167 Variovorax
14.3k
1.27k Alphaproteobacteria
306 Pasteurella multocida
1.37k Burkholderiales
Bacteria
1.75k Betaproteobacteria
320 Pasteurellaceae
306 Pasteurella
79 Escherichia coli
9.84k
1.00k Enterobacterales
652 Enterobacteriaceae
52 Salmonella enterica
Proteobacteria
226 Moraxellaceae
168 Acinetobacter
7.60k
7.37k
7.33k
435 Pseudomonas fluorescens
Gammaproteobacteria
Pseudomonadales
Pseudomonadaceae
Pseudomonas
86 Pseudomonas sp. MYb193
64 Viruses
16 Verrucomicrobia
D
P
C
O
F
G
S

**Fig. S8**

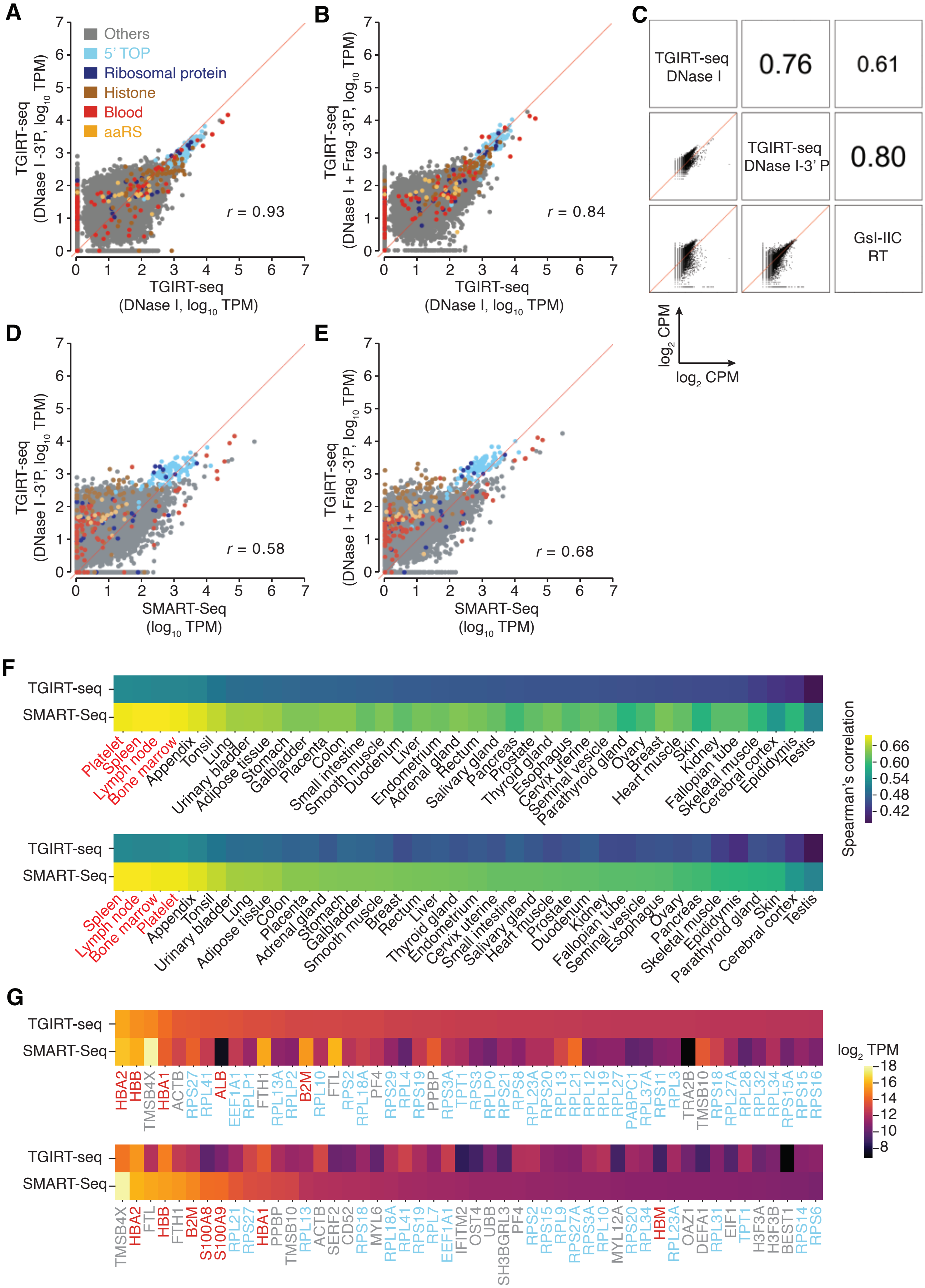
A
B
C
D
E
F
G
Others
5' TOP
Ribosomal protein
Histone
Blood
aaRS
TGIRT-seq (DNase I -3'P, log10 TPM)
TGIRT-seq (DNase I, log10 TPM)
r = 0.93
TGIRT-seq (DNase I + Frag -3'P, log10 TPM)
r = 0.84
TGIRT-seq DNase I
0.76
0.61
TGIRT-seq DNase I-3' P
0.80
GsI-IIC RT
log2 CPM
log2 CPM
SMART-Seq (log10 TPM)
r = 0.58
r = 0.68
TGIRT-seq
SMART-Seq
Spearman's correlation
0.66
0.60
0.54
0.48
0.42
Platelet
Spleen
Lymph node
Bone marrow
Appendix
Tonsil
Lung
Urinary bladder
Adipose tissue
Stomach
Galbladder
Placenta
Colon
Small intestine
Smooth muscle
Duodenum
Liver
Endometrium
Adrenal gland
Rectum
Salivary gland
Pancreas
Prostate
Thyroid gland
Esophagus
Cervix uterine
Seminal vesicle
Parathyroid gland
Ovary
Breast
Heart muscle
Skin
Kidney
Fallopian tube
Skeletal muscle
Cerebral cortex
Epididymis
Testis
log2 TPM
18
16
14
12
10
8
HBA2 HBB TMSB4X HBA1 ACTB RPS27 RPL41 ALB EEF1A1 RPLP1 FTH1 RPL13A RPLP2 B2M RPL10 FTL RPS2 RPL18A PF4 RPS29 RPL4 RPS19 PPBP RPS3A TPT1 RPS3 RPLP0 RPS21 RPS8 RPL23A RPS20 RPL13 RPL21 RPL12 RPL19 RPL27 PABPC1 RPL37A RPS11 RPL3 TRA2B TMSB10 RPS18 RPL27A RPL28 RPL32 RPL34 RPS15A RPS15 RPS16
TMSB4X HBA2 FTL HBB FTH1 B2M S100A8 S100A9 RPL21 RPS27 HBA1 PPBP TMSB10 RPL13 ACTB SERF2 CD52 RPS18 MYL6 RPL18A RPL41 RPS19 RPL7 EEF1A1 IFITM2 OST4 UBB SH3BGRL3 PF4 RPS2 RPS15 RPL9 RPS27A RPS3A RPL10 MYL12A RPS20 RPL34 HBM RPL23A OAZ1 DEFA1 RPL31 EIF1 TPT1 H3F3A H3F3B BEST1 RPS14 RPS6

### Fig. S9

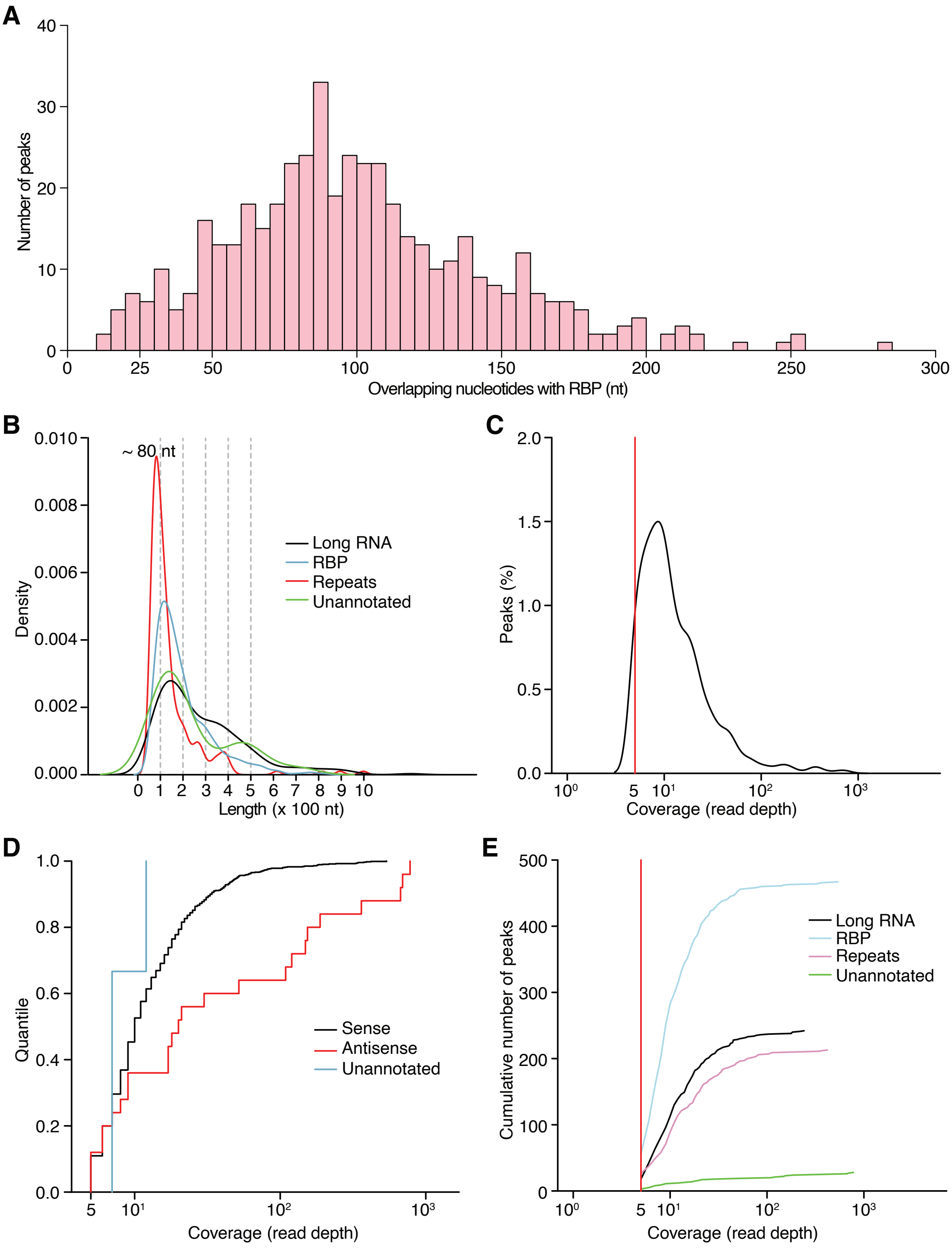
A
Number of peaks
Overlapping nucleotides with RBP (nt)
B
~ 80 nt
Density
Long RNA
RBP
Repeats
Unannotated
Length (x 100 nt)
C
Peaks (%)
Coverage (read depth)
D
Quantile
Sense
Antisense
Unannotated
Coverage (read depth)
E
Cumulative number of peaks
Long RNA
RBP
Repeats
Unannotated
Coverage (read depth)

### Fig. S10

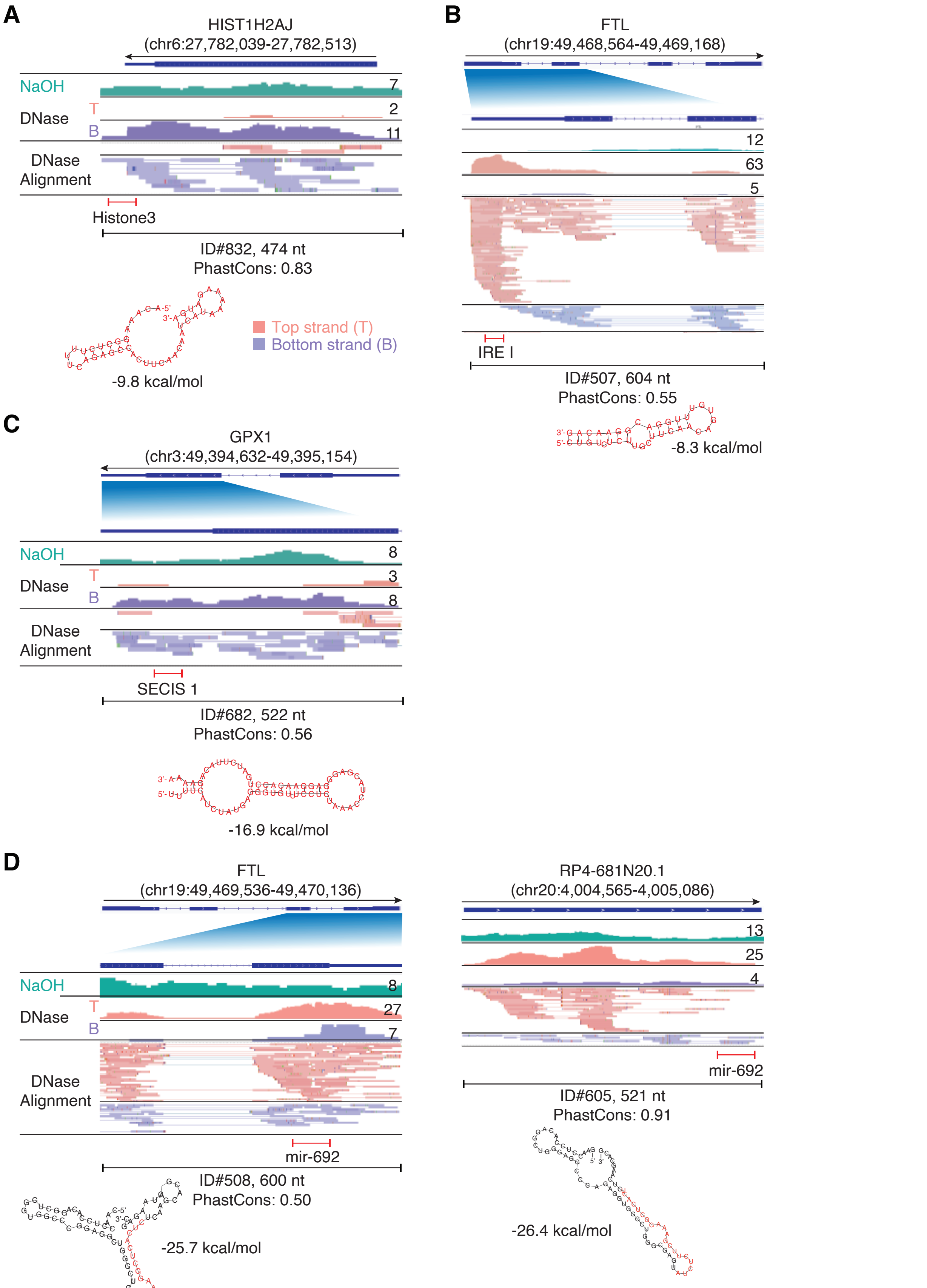


miR-692: CAACCAGCGCAGACUGGCGCGCCCCAGGGA–UCUCUGGGUGAGUAUCUCUUUGAGCGCCUCACUCUCAAGCACAACUAGGAGG

FTL: CAACCUCCACAGGCUGGGUGGCCCGGAGGCUGGGCUGGGCGAGUAUCUCUUCGAAAGGCUCACUCUCAAGCACGACUAAGAGC

RP4-681N20.1: GAACCUCCACAGGCUGGGAGGCCCAGAGG–UGGGCUGGGCGAGUAUCUCUUCGAAAGGCUCACUCUCAAGCACG

mature miR-692

### Fig. S11

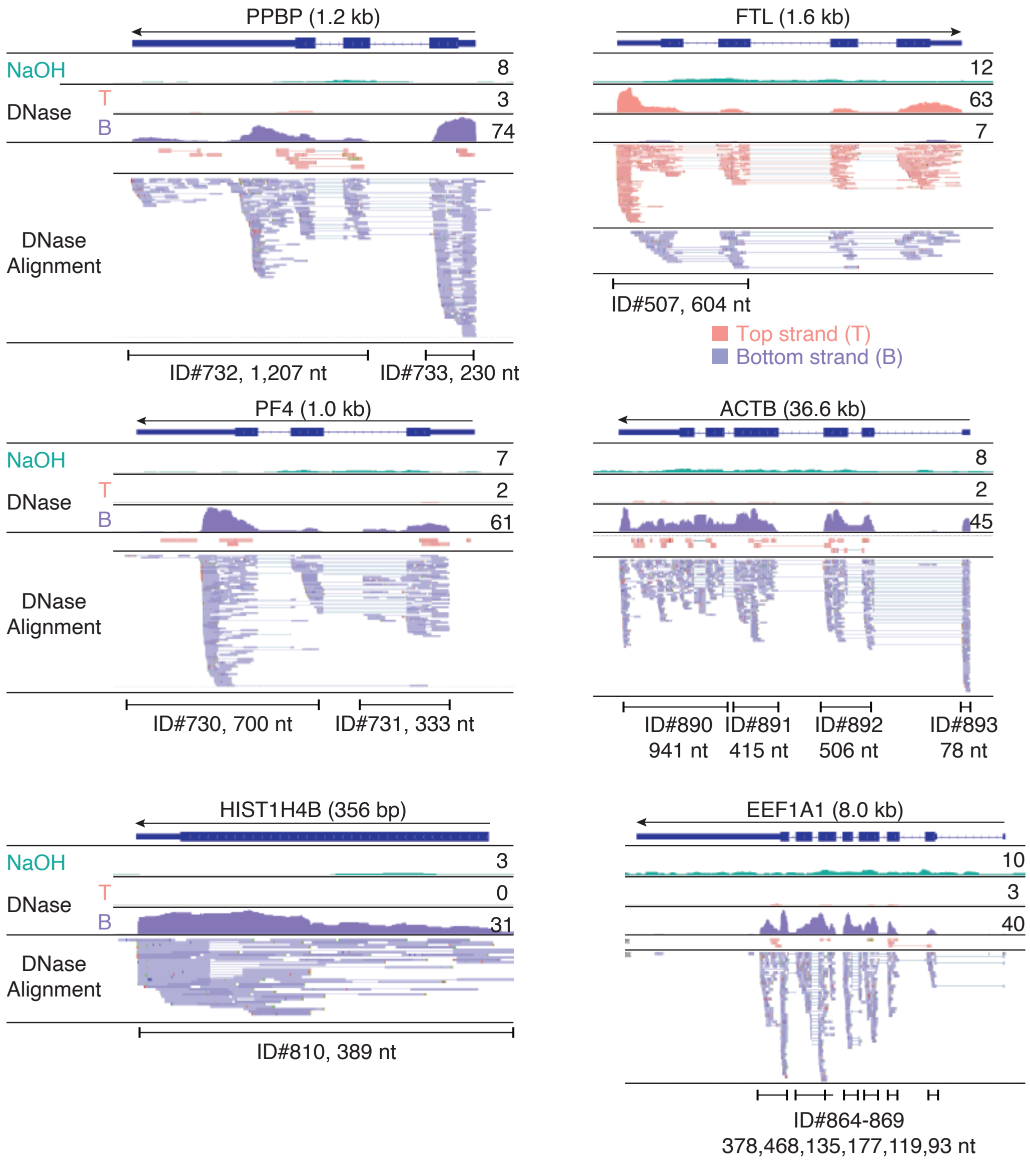
PPBP (1.2 kb)
NaOH
DNase
T
B
8
3
74
DNase
Alignment
ID#732, 1,207 nt
ID#733, 230 nt
FTL (1.6 kb)
12
63
7
ID#507, 604 nt
Top strand (T)
Bottom strand (B)
PF4 (1.0 kb)
7
2
61
ID#730, 700 nt
ID#731, 333 nt
ACTB (36.6 kb)
8
2
45
ID#890
941 nt
ID#891
415 nt
ID#892
506 nt
ID#893
78 nt
HIST1H4B (356 bp)
3
0
31
ID#810, 389 nt
EEF1A1 (8.0 kb)
10
3
40
ID#864-869
378,468,135,177,119,93 nt

#### Fig. S12

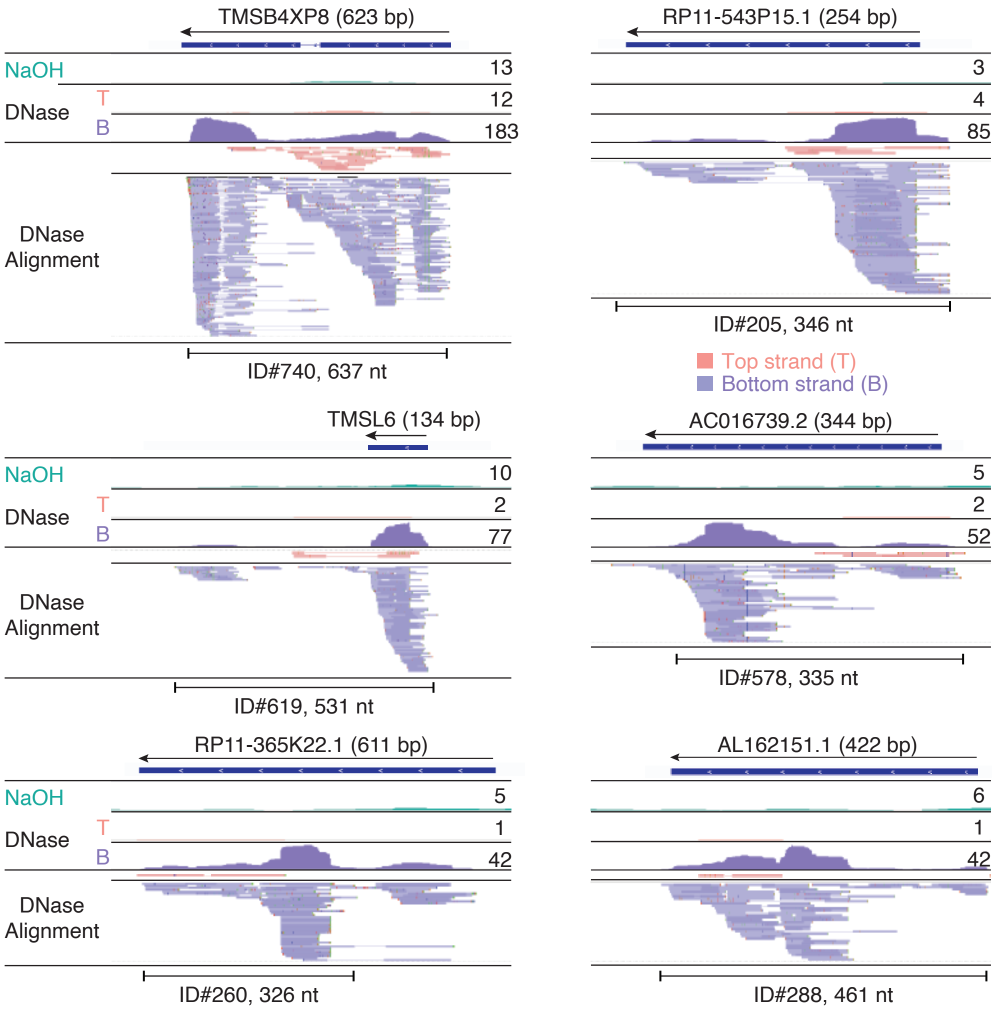

TMSB4XP8 (623 bp)
NaOH
DNase
T
B
13
12
183
DNase Alignment
ID#740, 637 nt
RP11-543P15.1 (254 bp)
3
4
85
ID#205, 346 nt
Top strand (T)
Bottom strand (B)
TMSL6 (134 bp)
NaOH
DNase
T
B
10
2
77
DNase Alignment
ID#619, 531 nt
AC016739.2 (344 bp)
5
2
52
ID#578, 335 nt
RP11-365K22.1 (611 bp)
NaOH
DNase
T
B
5
1
42
DNase Alignment
ID#260, 326 nt
AL162151.1 (422 bp)
6
1
42
ID#288, 461 nt

### Fig. S13

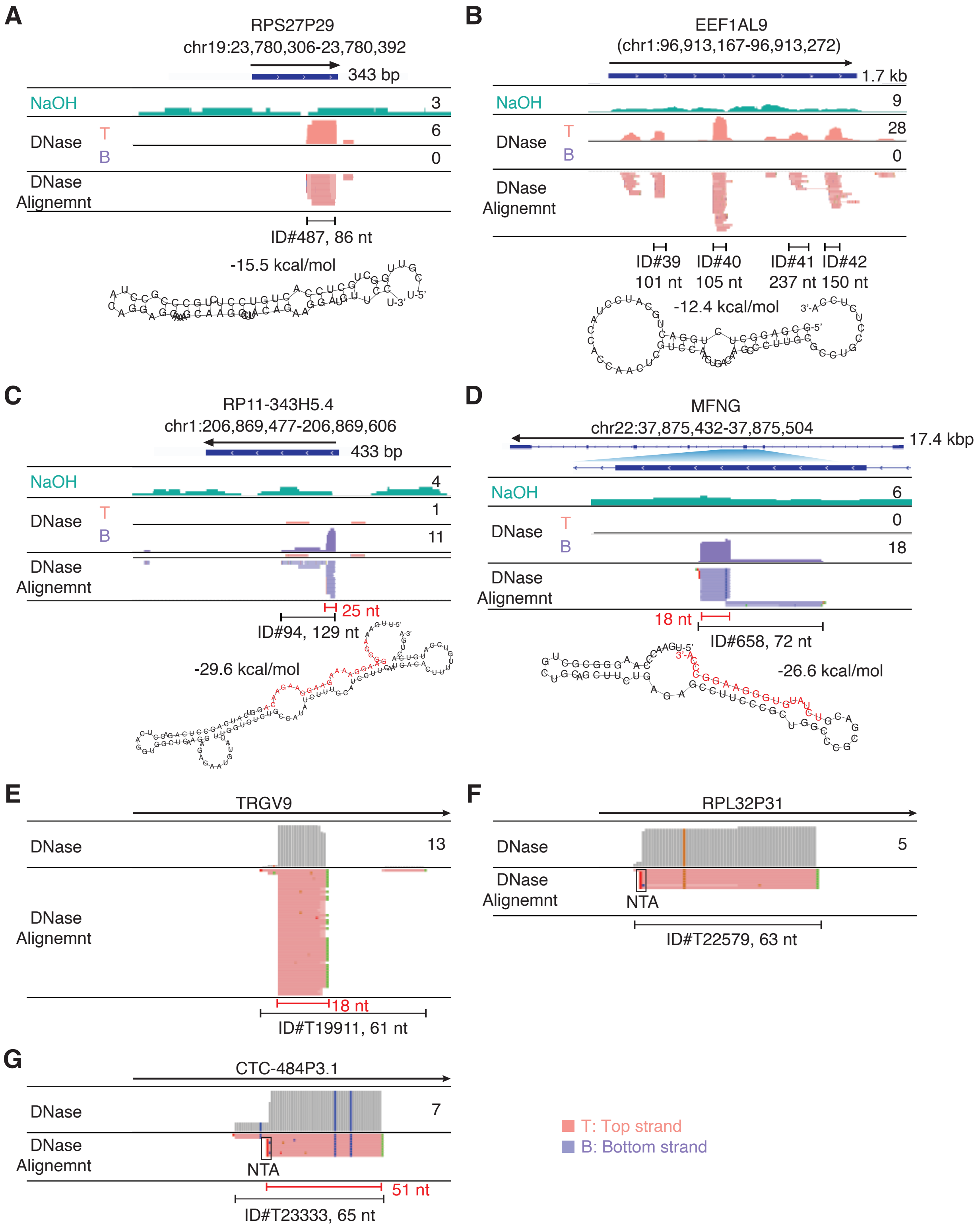
A
RPS27P29
chr19:23,780,306-23,780,392
343 bp
NaOH
3
DNase
T
B
6
0
DNase Alignemnt
ID#487, 86 nt
-15.5 kcal/mol
B
EEF1AL9
(chr1:96,913,167-96,913,272)
1.7 kb
NaOH
9
DNase
T
B
28
0
DNase Alignemnt
ID#39 101 nt
ID#40 105 nt
ID#41 237 nt
ID#42 150 nt
-12.4 kcal/mol
C
RP11-343H5.4
chr1:206,869,477-206,869,606
433 bp
NaOH
4
DNase
T
B
1
11
DNase Alignemnt
25 nt
ID#94, 129 nt
-29.6 kcal/mol
D
MFNG
chr22:37,875,432-37,875,504
17.4 kbp
NaOH
6
DNase
T
B
0
18
DNase Alignemnt
18 nt
ID#658, 72 nt
-26.6 kcal/mol
E
TRGV9
DNase
13
DNase Alignemnt
18 nt
ID#T19911, 61 nt
F
RPL32P31
DNase
5
DNase Alignemnt
NTA
ID#T22579, 63 nt
G
CTC-484P3.1
DNase
7
DNase Alignemnt
NTA
51 nt
ID#T23333, 65 nt
T: Top strand
B: Bottom strand

Fig. S14

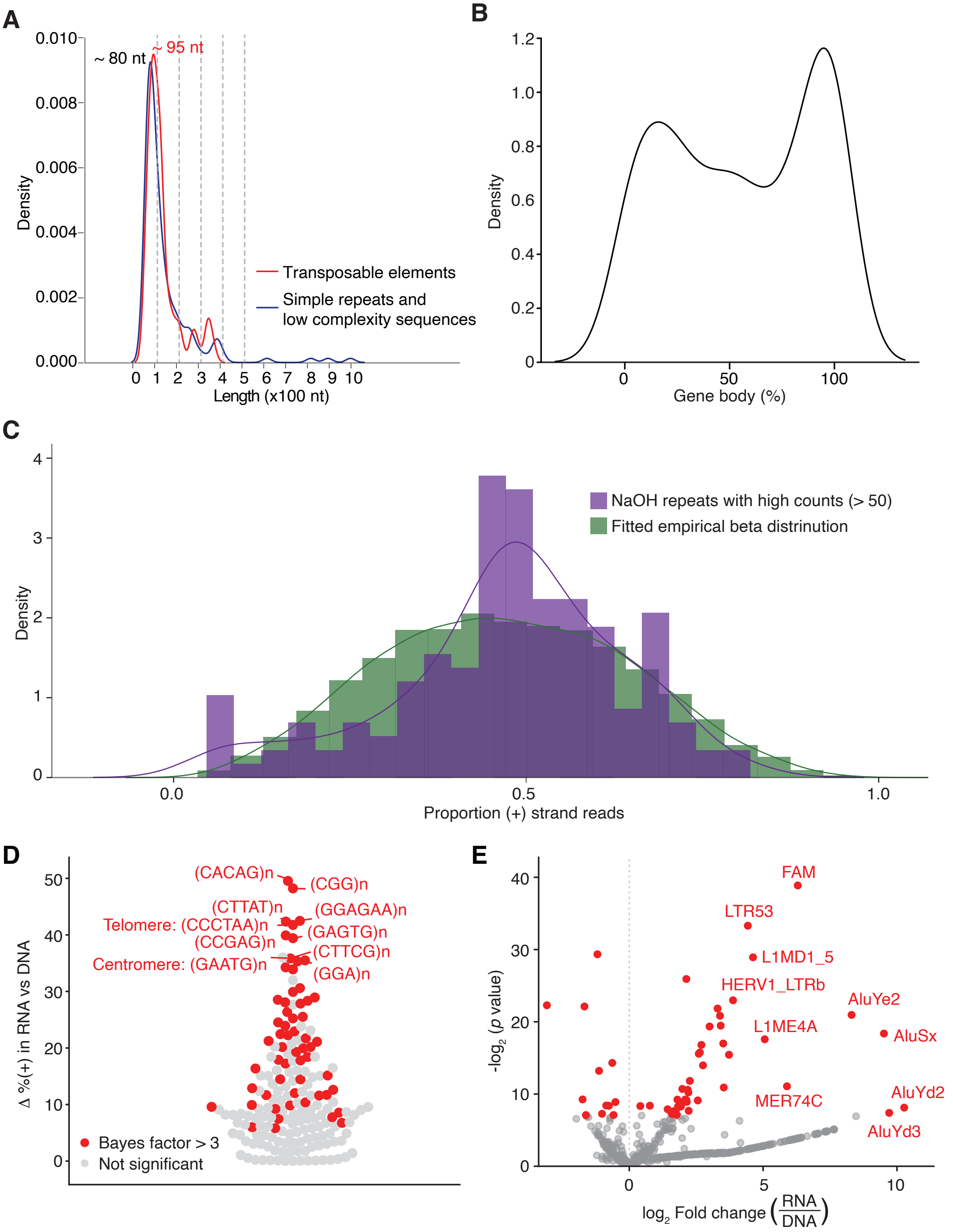
A
~ 80 nt
~ 95 nt
Density
Length (x100 nt)
Transposable elements
Simple repeats and
low complexity sequences
B
Density
Gene body (%)
C
NaOH repeats with high counts (> 50)
Fitted empirical beta distrinution
Density
Proportion (+) strand reads
D
(CACAG)n
(CGG)n
(CTTAT)n
(GGAGAA)n
Telomere: (CCCTAA)n
(GAGTG)n
(CCGAG)n
(CTTCG)n
Centromere: (GAATG)n
(GGA)n
Δ %(+) in RNA vs DNA
Bayes factor > 3
Not significant
E
FAM
LTR53
L1MD1_5
HERV1_LTRb
AluYe2
AluSx
L1ME4A
MER74C
AluYd2
AluYd3
-log2 (p value)
log2 Fold change (RNA/DNA)

### Fig. S15

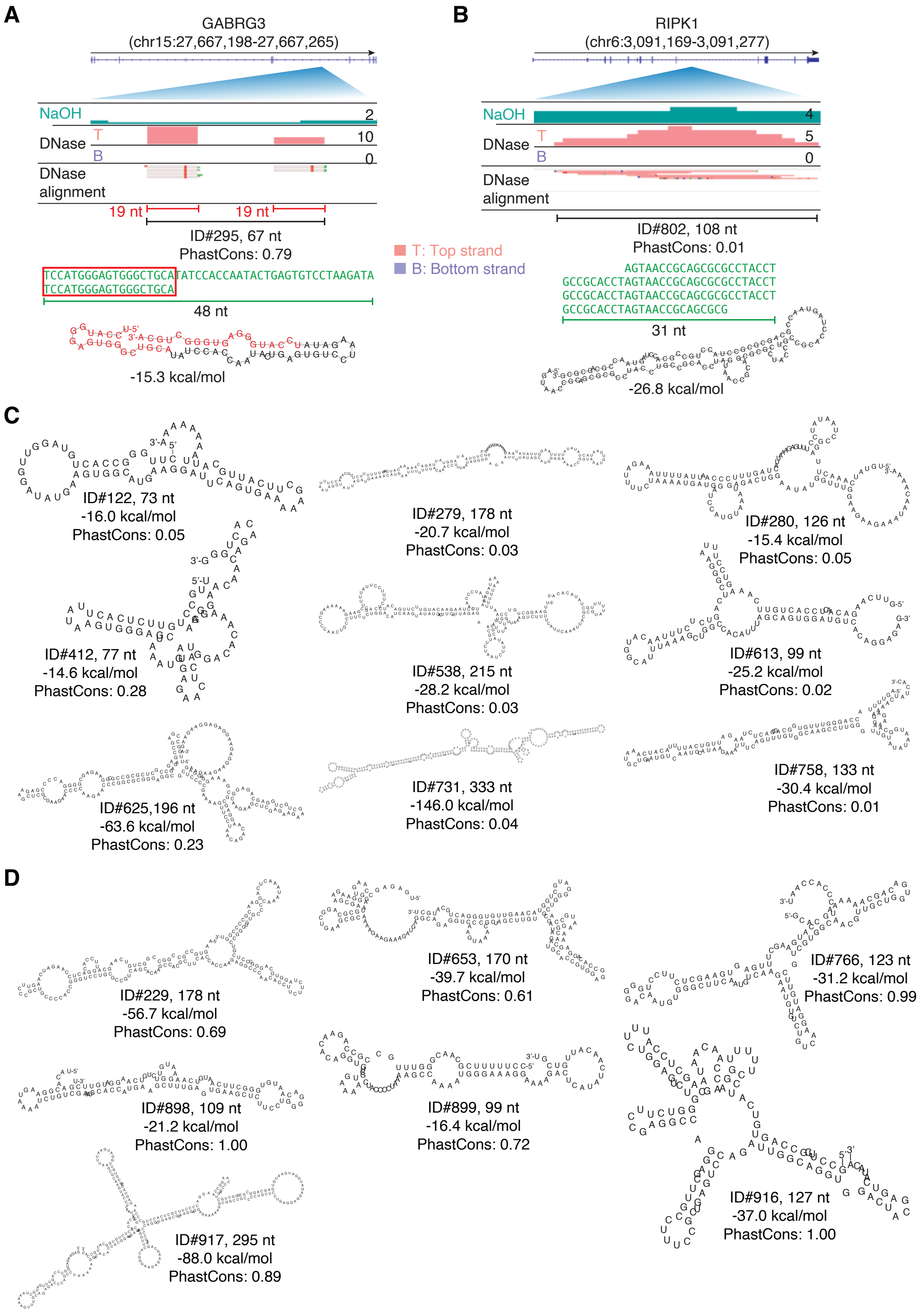
A
GABRG3
(chr15:27,667,198-27,667,265)
NaOH
2
DNase
T
10
B
0
DNase alignment
19 nt
19 nt
ID#295, 67 nt
PhastCons: 0.79
TCCATGGGAGTGGGCTGCATATCCACCAATACTGAGTGTCCTAAGATA
TCCATGGGAGTGGGCTGCA
48 nt
-15.3 kcal/mol
T: Top strand
B: Bottom strand
B
RIPK1
(chr6:3,091,169-3,091,277)
NaOH
4
DNase
T
5
B
0
DNase alignment
ID#802, 108 nt
PhastCons: 0.01
AGTAACCGCAGCGCGCCTACCT
GCCGCACCTAGTAACCGCAGCGCGCCTACCT
GCCGCACCTAGTAACCGCAGCGCGCCTACCT
GCCGCACCTAGTAACCGCAGCGCG
31 nt
-26.8 kcal/mol
C
ID#122, 73 nt
-16.0 kcal/mol
PhastCons: 0.05
ID#279, 178 nt
-20.7 kcal/mol
PhastCons: 0.03
ID#280, 126 nt
-15.4 kcal/mol
PhastCons: 0.05
ID#412, 77 nt
-14.6 kcal/mol
PhastCons: 0.28
ID#538, 215 nt
-28.2 kcal/mol
PhastCons: 0.03
ID#613, 99 nt
-25.2 kcal/mol
PhastCons: 0.02
ID#625,196 nt
-63.6 kcal/mol
PhastCons: 0.23
ID#731, 333 nt
-146.0 kcal/mol
PhastCons: 0.04
ID#758, 133 nt
-30.4 kcal/mol
PhastCons: 0.01
D
ID#229, 178 nt
-56.7 kcal/mol
PhastCons: 0.69
ID#653, 170 nt
-39.7 kcal/mol
PhastCons: 0.61
ID#766, 123 nt
-31.2 kcal/mol
PhastCons: 0.99
ID#898, 109 nt
-21.2 kcal/mol
PhastCons: 1.00
ID#899, 99 nt
-16.4 kcal/mol
PhastCons: 0.72
ID#916, 127 nt
-37.0 kcal/mol
PhastCons: 1.00
ID#917, 295 nt
-88.0 kcal/mol
PhastCons: 0.89

#### Fig. S16

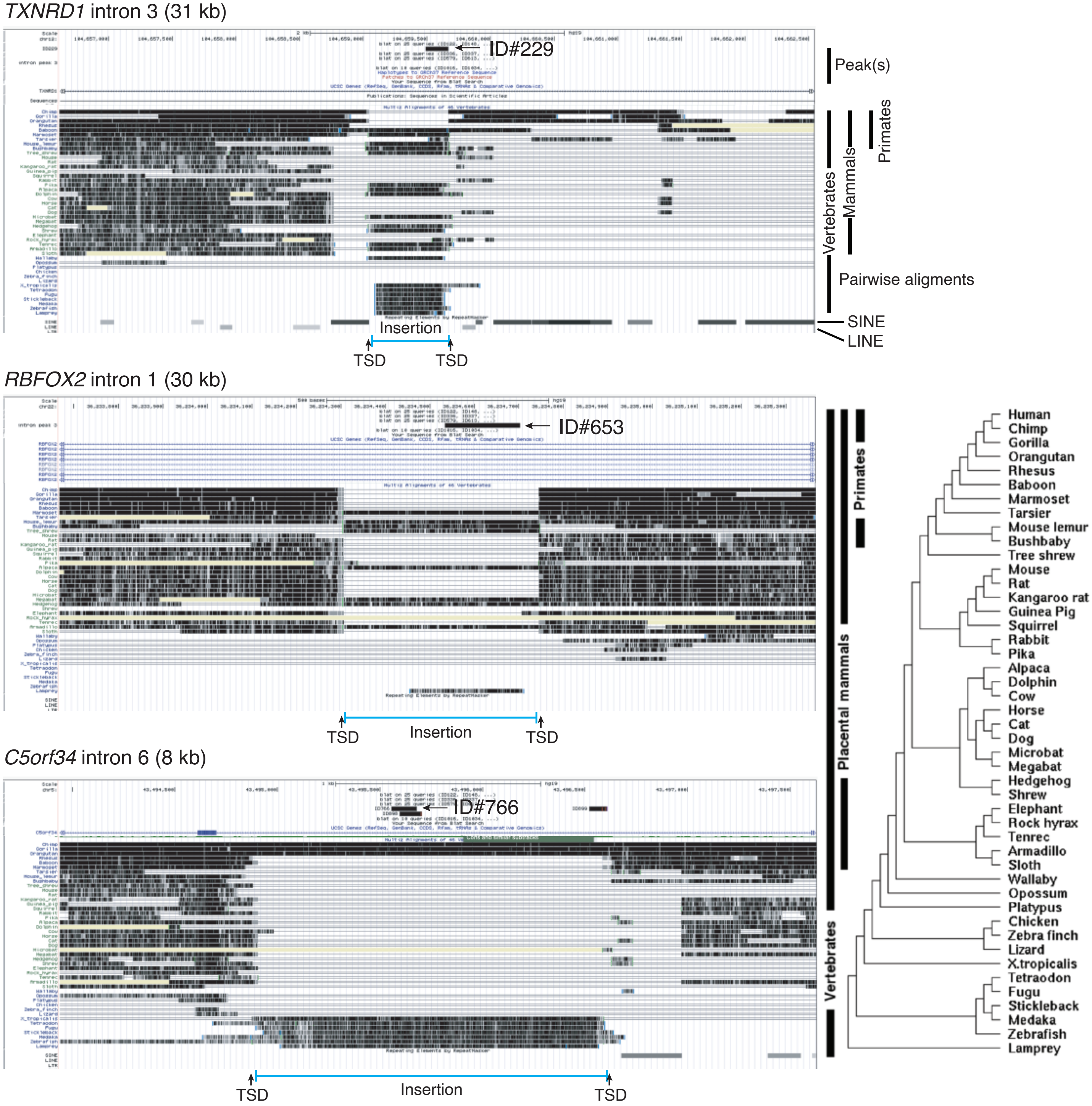

TXNRD1 intron 3 (31 kb)
ID#229
Peak(s)
Primates
Mammals
Vertebrates
Pairwise aligments
SINE
LINE
Insertion
TSD
TSD
RBFOX2 intron 1 (30 kb)
ID#653
TSD
Insertion
TSD
C5orf34 intron 6 (8 kb)
ID#766
TSD
Insertion
TSD
Primates
Placental mammals
Vertebrates
Human
Chimp
Gorilla
Orangutan
Rhesus
Baboon
Marmoset
Tarsier
Mouse lemur
Bushbaby
Tree shrew
Mouse
Rat
Kangaroo rat
Guinea Pig
Squirrel
Rabbit
Pika
Alpaca
Dolphin
Cow
Horse
Cat
Dog
Microbat
Megabat
Hedgehog
Shrew
Elephant
Rock hyrax
Tenrec
Armadillo
Sloth
Wallaby
Opossum
Platypus
Chicken
Zebra finch
Lizard
X.tropicalis
Tetraodon
Fugu
Stickleback
Medaka
Zebrafish
Lamprey

#### Fig. S16 (continued)

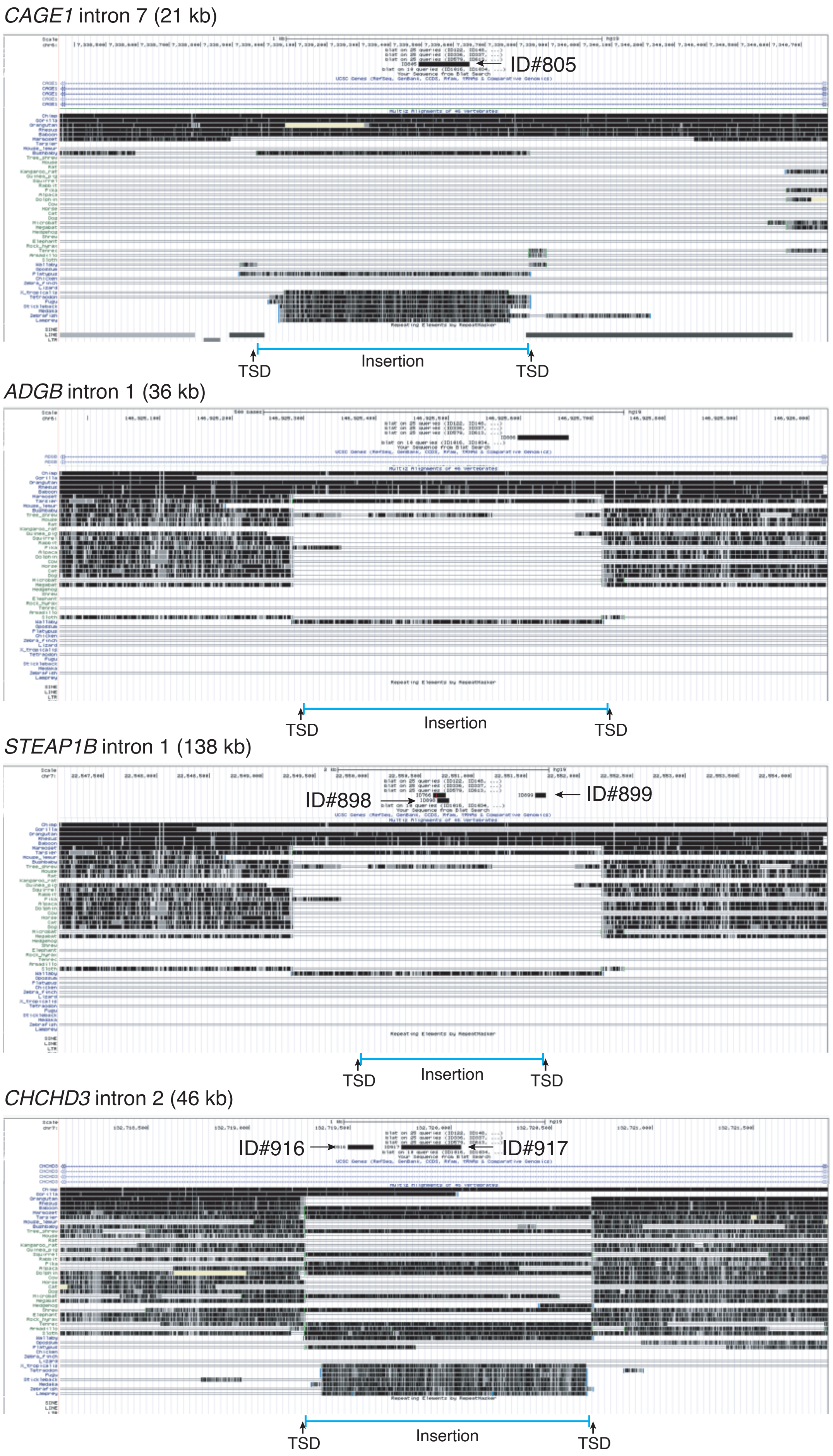

**Fig. S17**

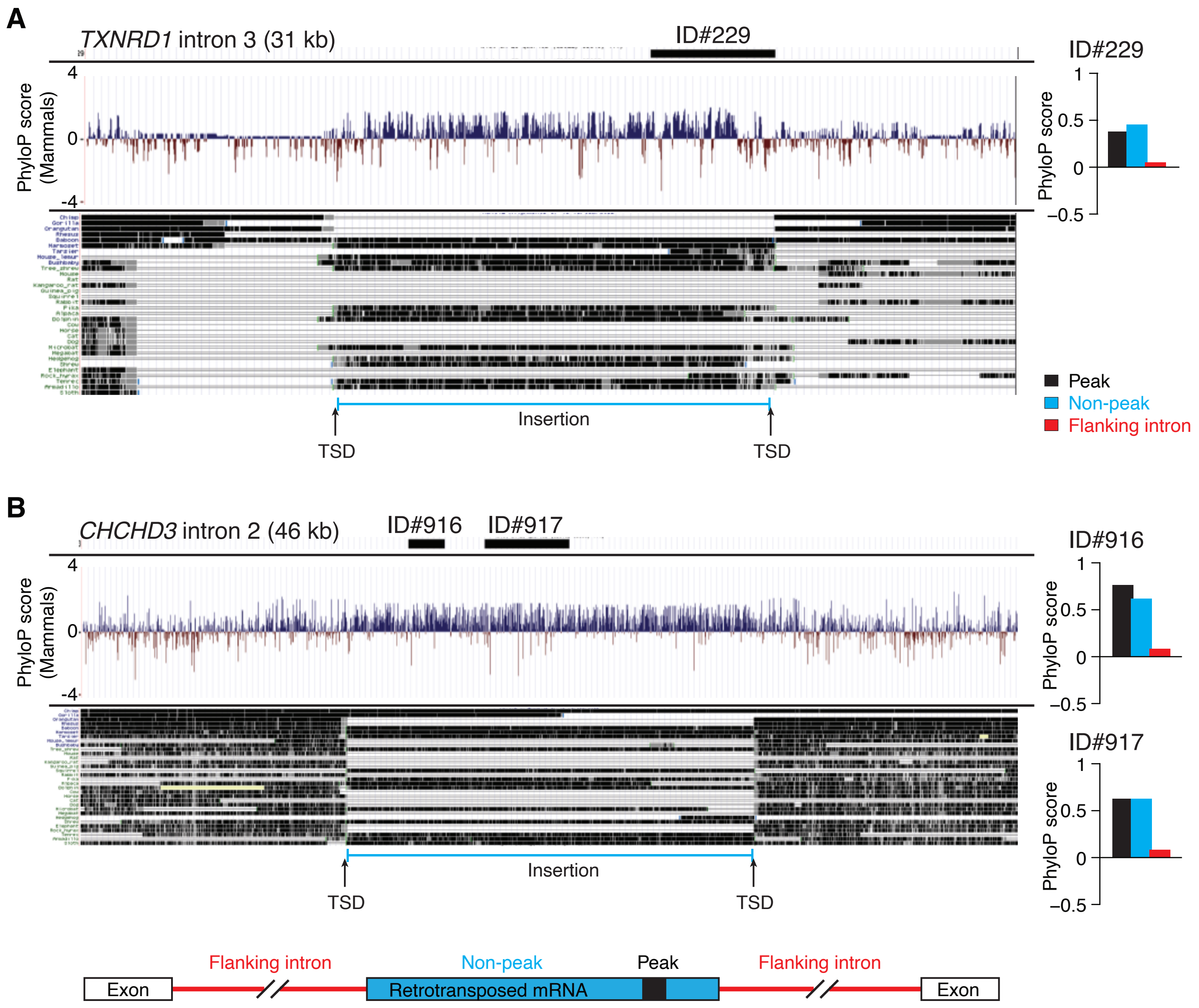

A
TXNRD1 intron 3 (31 kb)
ID#229
PhyloP score (Mammals)
4
0
-4
Insertion
TSD
TSD
ID#229
PhyloP score
1
0.5
0
−0.5
Peak
Non-peak
Flanking intron
B
CHCHD3 intron 2 (46 kb)
ID#916
ID#917
PhyloP score (Mammals)
4
0
-4
Insertion
TSD
TSD
ID#916
PhyloP score
1
0.5
0
−0.5
ID#917
PhyloP score
1
0.5
0
−0.5
Flanking intron
Non-peak
Peak
Flanking intron
Exon
Retrotransposed mRNA
Exon

### Fig. S18

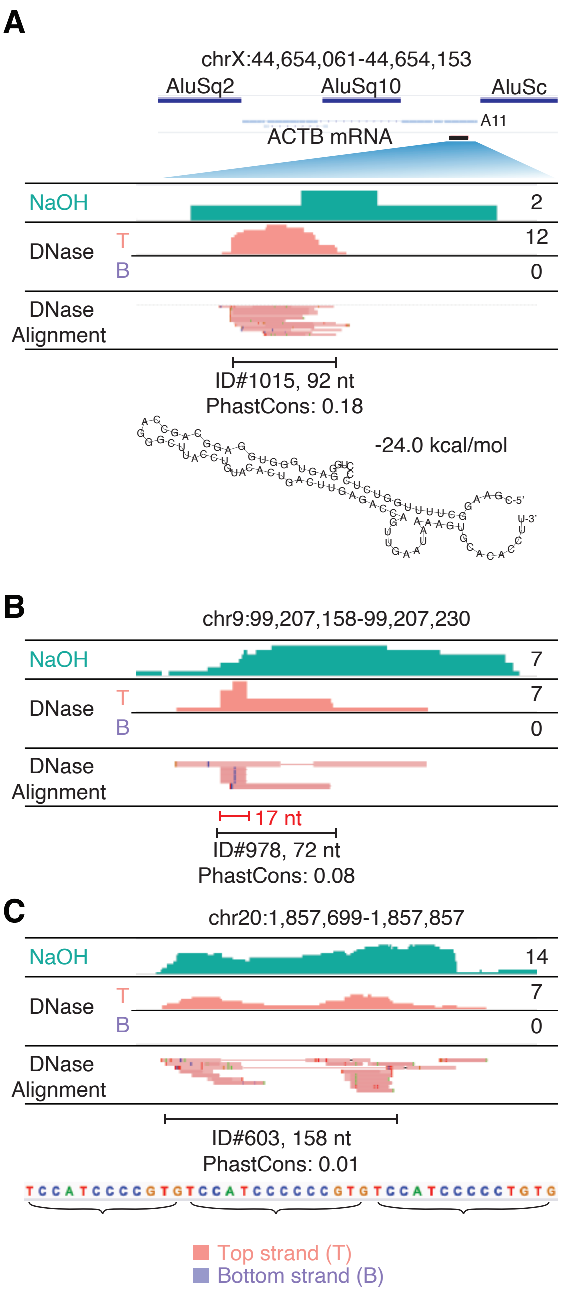

A
chrX:44,654,061-44,654,153
AluSq2
AluSq10
AluSc
A11
ACTB mRNA
NaOH
2
DNase
T
B
12
0
DNase
Alignment
ID#1015, 92 nt
PhastCons: 0.18
-24.0 kcal/mol
B
chr9:99,207,158-99,207,230
NaOH
7
DNase
T
B
7
0
DNase
Alignment
17 nt
ID#978, 72 nt
PhastCons: 0.08
C
chr20:1,857,699-1,857,857
NaOH
14
DNase
T
B
7
0
DNase
Alignment
ID#603, 158 nt
PhastCons: 0.01
TCCATCCCCGTGTCCATCCCCCCGTGTCCATCCCCCTGTG
Top strand (T)
Bottom strand (B)

#### Fig. S19

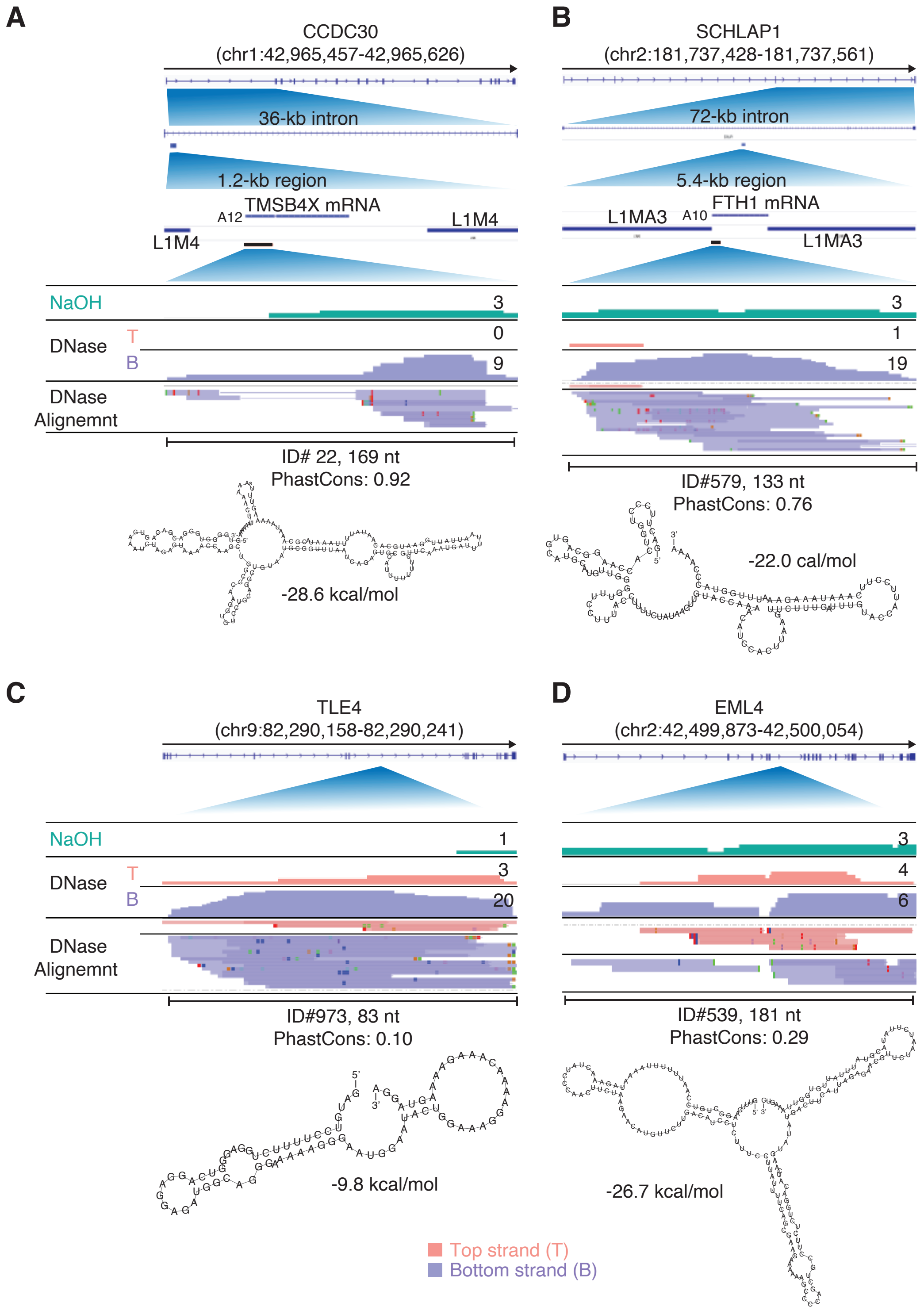

A
CCDC30
(chr1:42,965,457-42,965,626)
36-kb intron
1.2-kb region
A12
TMSB4X mRNA
L1M4
L1M4
NaOH
3
DNase
T
0
B
9
DNase Alignemnt
ID# 22, 169 nt
PhastCons: 0.92
-28.6 kcal/mol
B
SCHLAP1
(chr2:181,737,428-181,737,561)
72-kb intron
5.4-kb region
L1MA3
A10
FTH1 mRNA
L1MA3
3
1
19
ID#579, 133 nt
PhastCons: 0.76
-22.0 cal/mol
C
TLE4
(chr9:82,290,158-82,290,241)
NaOH
1
DNase
T
3
B
20
DNase Alignemnt
ID#973, 83 nt
PhastCons: 0.10
-9.8 kcal/mol
D
EML4
(chr2:42,499,873-42,500,054)
3
4
6
ID#539, 181 nt
PhastCons: 0.29
-26.7 kcal/mol
Top strand (T)
Bottom strand (B)

**Fig. S20**

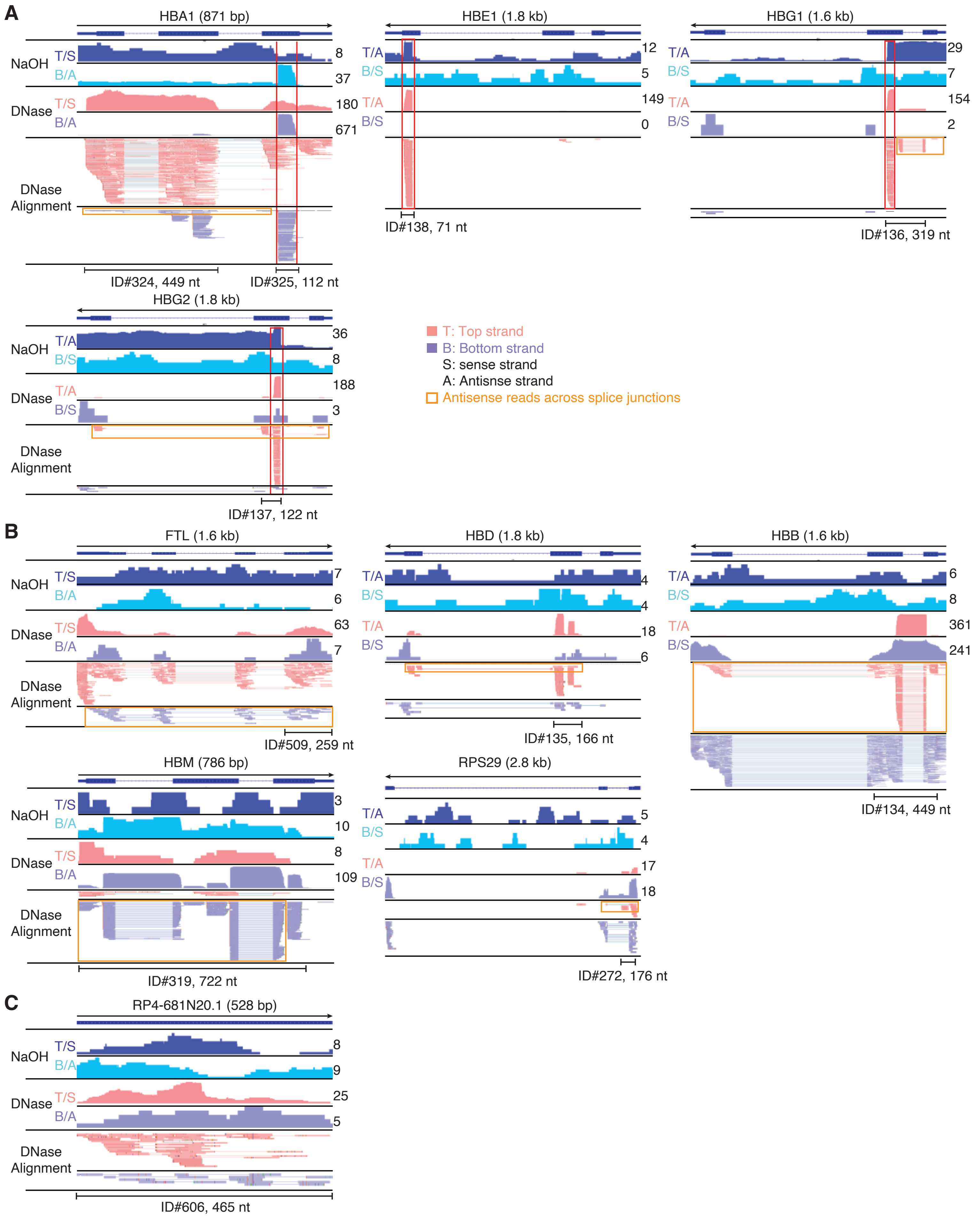

A
HBA1 (871 bp)
NaOH
T/S
B/A
8
37
DNase
T/S
B/A
180
671
DNase Alignment
ID#324, 449 nt
ID#325, 112 nt
HBE1 (1.8 kb)
T/A
B/S
12
5
149
0
ID#138, 71 nt
HBG1 (1.6 kb)
29
7
154
2
ID#136, 319 nt
HBG2 (1.8 kb)
36
8
188
3
ID#137, 122 nt
T: Top strand
B: Bottom strand
S: sense strand
A: Antisnse strand
Antisense reads across splice junctions
B
FTL (1.6 kb)
7
6
63
7
ID#509, 259 nt
HBD (1.8 kb)
4
4
18
6
ID#135, 166 nt
HBB (1.6 kb)
6
8
361
241
ID#134, 449 nt
HBM (786 bp)
3
10
8
109
ID#319, 722 nt
RPS29 (2.8 kb)
5
4
17
18
ID#272, 176 nt
C
RP4-681N20.1 (528 bp)
8
9
25
5
ID#606, 465 nt
